# Modulation of *Ideonella sakaiensis* PETase active site flexibility and activity on morphologically distinct substrates by surface charge engineering

**DOI:** 10.1101/2024.05.07.589851

**Authors:** Ke Ding, Zarina Levitskaya, Barindra Sana, Rupali Reddy Pasula, Srinivasaraghavan Kannan, Abdurrahman Adam, Vishnu Vadanan Sundaravadanam, Chandra Verma, Sierin Lim, John F. Ghadessy

## Abstract

Enzymatic hydrolysis of polyethylene terephthalate (PET) waste is a compelling strategy for environmentally friendly recycling of a major pollutant. Here, we investigate the effects of surface charge point mutations both proximal and distal to the active site of the mesophilic PET-degrading enzyme from *Ideonella sakaienses* (*Is*PETase) and an engineered thermostable variant with superior activity, STAR PETase. The vicinal K95A mutation significantly inhibited *Is*PETase activity on mechanically processed PET powder. Conversely, this mutation significantly increased hydrolysis of PET powder in the STAR PETase background. Activity of both enzymes on PET film was inhibited by the K95A mutation, highlighting complex interplay between mutation context and substrate morphology. Further installing the distal R132N and R280A surface charge mutations potentiated activity of STAR on all substrates tested. This variant afforded 100% degradation of bottle-grade PET powder in 3 days at 40°C reaction temperature, a 3-fold improvement over *Is*PETase. Molecular dynamics simulations reveal modulation of active site flexibility in mutants, which differentially impacts both hydrolysis of morphologically distinct PET substrates and the concentration-dependent inhibition phenomenon observed for PETase.

## Introduction

Over the last 20 years scientists have described several microbial enzymes that can hydrolyse polyethylene terephthalate (PET), one of the most abundant plastics materials on earth widely used in disposable food and drink containers^1-3^. Particular interest lies in exploiting these enzymes to enable re/up-cycling of discarded and accumulating PET waste, a significant global issue^4^. Endogenous enzymes described thus far show sub-optimal activities on highly crystalline commercial PET^5-7^, mandating protein engineering to boost performance to economically viable levels^8-10^. To date, an engineered variant of leaf branch compost (LCC) cutinase is the only enzyme being tested at pilot scale for recycling of PET waste mechanically pre-treated to reduce crystallinity^11^. *Is*PETase (hereafter referred to as PETase) elaborated by the bacterium *Ideonella sakaiensis* has also garnered much attention^2^. Numerous variants with improved activity and robustness have been developed by employing various engineering strategies^12-22^. Computational design and machine learning approaches were respectively used to derive the more efficient and thermostable Dura- and FAST PETase variants^8,9^. Random mutagenesis and screening have yielded the highly active Depo- and HOT PETase variants^5,23^. Rational structure-guided engineering has also generated PETase iterations with notable increases in thermostability, activity, solubility, and resistance to inhibition^6,24,25^.

Analysis of both natural and synthetic enzyme phylogenies suggests a relationship between surface charge distribution and PET hydrolysis activity^26-28^. Both the PETase substrate binding groove and the enzyme face presenting it are notably cationic, an evolutionary adaption differentiating it from most phylogenetically related enzymes (cutinases, lipases, carboxylesterases) that show reduced or absent PET degradation activity^12,26^. Along with adaptations rendering its active site more flexible^29^, surface charge likely plays an important role in mediating productive interactions at the liquid-solid interface of recalcitrant crystalline PET substrates^30^. Pre-treatment of PET with anionic surfactants can accelerate enzymatic PET degradation by up to 120-fold^31^, highlighting both the importance of electrostatic interactions for substrate binding and the potential to further improve through engineering. Accordingly, mutation of charged surface residues is observed in many described PETase variants with improved PET hydrolytic activity. Generally, removal of positive charged residues either proximal or co-facial to the substrate binding pocket reduces activity on negatively charged PET substrate^31,32^. Conversely, loss of positive charge at residues distal to the active site is observed in several catalytically improved variants^5,9,25^. As seen in other proteins^33^, surface charge modifications can also increase thermostability of PET hydrolases, likely contributing to the associated activity enhancements over longer term (days) incubation periods.

In this study we examined the effects of two surface charge mutations, one proximal and the other distal, to the PETase substrate binding channel. Mutations were introduced into wild type PETase and an engineered thermostable derivative. We observed that activity profiles of mutants could dramatically differ based on parental enzyme scaffold and PET substrate morphology i.e., mechanically pre-processed powder or film. Molecular dynamics simulations provide further insight into relationship between PETase active site flexibility and activity on different PET substrates.

## Results

### Selection of surface charged residues

To further understand the role of surface charge in PET hydrolysis we investigated mutation of charged residues both co-facial and distal to the PETase active site (Fig. 1). K95 resides ~15Å from the substrate binding pocket, and along with R90 and R53 forms a cationic surface patch co-facial to the PET binding pocket. R132 sits on the back face of the enzyme relative to the active site at a distance of ~22Å. At both these positions, conservation of positive charge is not absolute in evolutionary related cutinases/hydrolases, suggesting possible gain of function related to the superior PET degradation observed in PETase^14^. Both residues were investigated in the context of wild type PETase and a variant with improved thermostability and activity, STAR PETase. We designed the latter based on structural homology modeling with a thermostabilized and highly active cutinase variant^34^, with introduction of mutations Q119G/N233C/S238W/S282C (Fig. 1). This mutation set in PETase has also been described in a patent application by Carbios^35^. STAR PETase showed optimal activity at pH8.0 and 40°C incubation temperature on lcPET powder substrate (Fig. S1). Based on phylogenetic analysis and ancestral sequence reconstruction^14,26^, K95 and R132 were respectively mutated to alanine and asparagine, evolutionary favoured non-charged amino acids found at the corresponding positions in related enzymes (Fig. S2).

**Figure 1.**
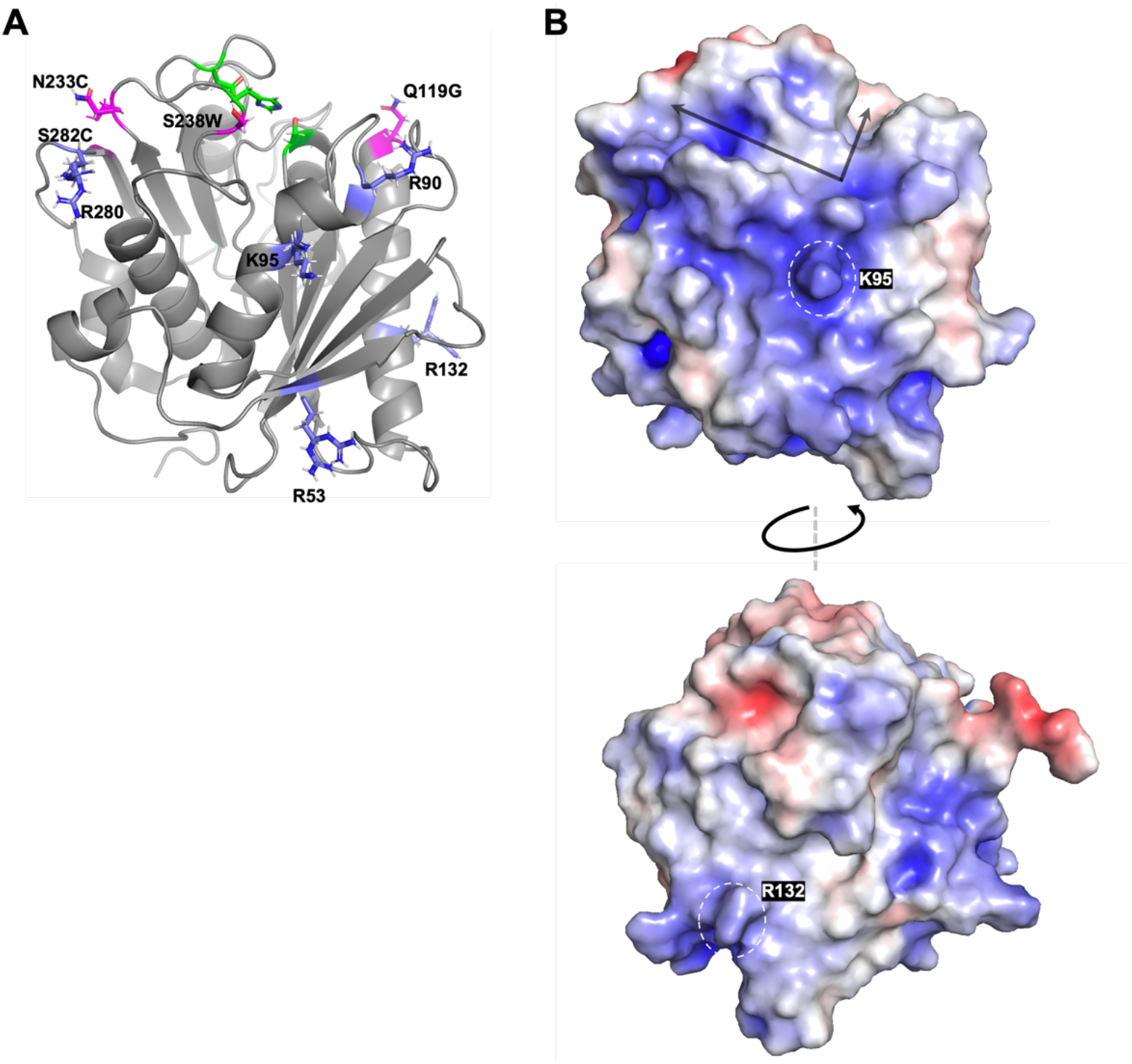
Structure of *Is*PETase. (A) Catalytic triad residues S160, D206, H237 are highlighted in green. Mutations in STAR PETase highlighted in magenta. Charged residues mutated in this study coloured slate. (B) *Is*PETase surface electrostatic potential with negative and positive regions depicted red and blue respectively. Active site cleft arrowed. Images generated from PDB:6eqe^36^ using PyMOL^65^.

### Effects of surface charge mutations on enzyme thermostabilities

Surface charge modifications can impart both positive and negative changes in protein stability^37^. We therefore measured thermostability of PETase and STAR enzymes with K95A and/or R132N mutations introduced using differential scanning fluorimetry^38^ (Table 1). STAR PETase showed a 14°C increased T_m_ (57.5°C) over PETase (43.5°C), with a significant contribution likely arising from a disulfide linkage introduced between C233 and C282^25^. The K95A mutation generally increased T_m_ by ~ 1-1.5°C across both PETase scaffolds, whilst R132N decreased T_m_ by ~ 0.5-1°C. These changes were effectively cancelled out in the double mutants which showed no significant change in T_m_. Further installation of the R280A surface charge mutation, previously shown to increase activity of PETase^14^ increased the T_m_s of STAR and STAR^K95A/R132N^ to 61.67°C and 61.25°C respectively.

**Table 1.**
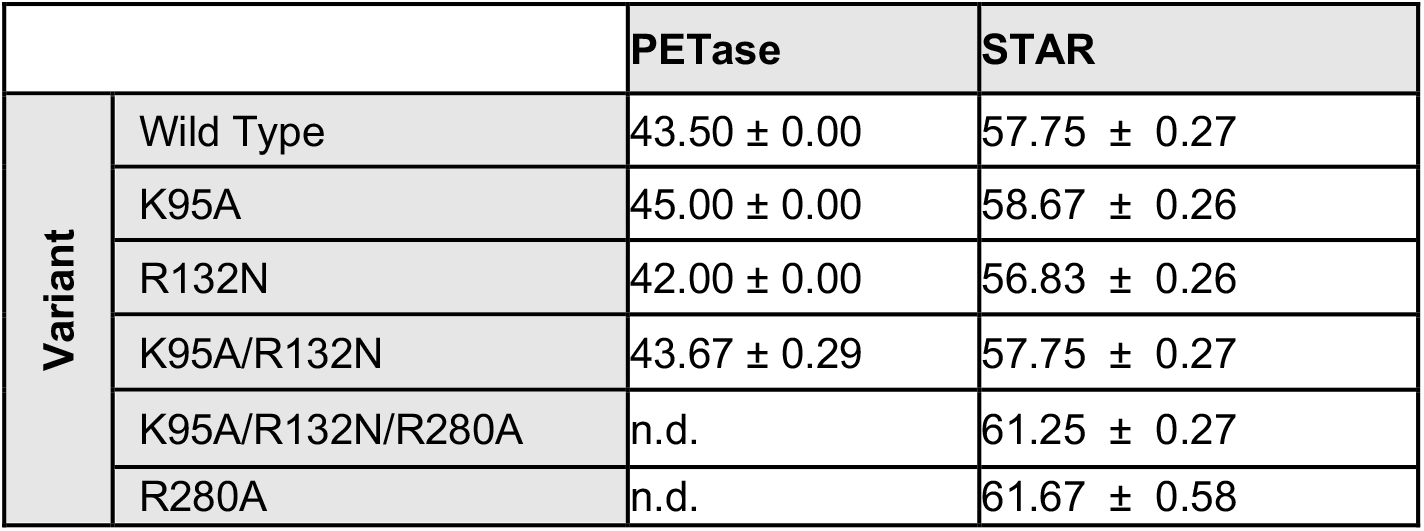
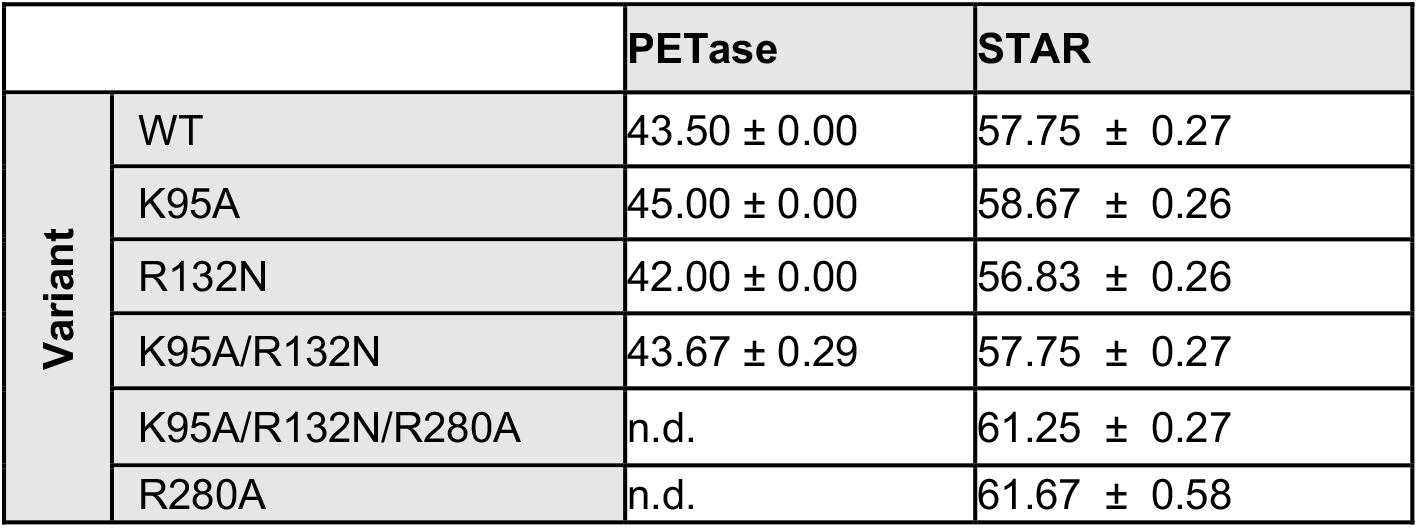
Melting temperatures of PETase, STAR PETase, and indicated variants (mean ± SD, n = 3-6)

|  |  | PETase | STAR |
| --- | --- | --- | --- |
| Variant | Wild Type | 43.50 ± 0.00 | 57.75 ± 0.27 |
|  | K95A | 45.00 ± 0.00 | 58.67 ± 0.26 |
|  | R132N | 42.00 ± 0.00 | 56.83 ± 0.26 |
|  | K95A/R132N | 43.67 ± 0.29 | 57.75 ± 0.27 |
|  | K95A/R132N/R280A | n.d. | 61.25 ± 0.27 |
|  | R280A | n.d. | 61.67 ± 0.58 |

|  |  | PETase | STAR |
| --- | --- | --- | --- |
| Variant | WT | 43.50 ± 0.00 | 57.75 ± 0.27 |
|  | K95A | 45.00 ± 0.00 | 58.67 ± 0.26 |
|  | R132N | 42.00 ± 0.00 | 56.83 ± 0.26 |
|  | K95A/R132N | 43.67 ± 0.29 | 57.75 ± 0.27 |
|  | K95A/R132N/R280A | n.d. | 61.25 ± 0.27 |
|  | R280A | n.d. | 61.67 ± 0.58 |

### Activity analysis of enzyme variants

Enzyme activity was next measured using micronized PET powder (8% crystallinity) and film substrates (Goodfellow lc1445, 7% crystallinity) with respective enzyme loadings (W_enzyme_/W_PET)_ of 0.13% and 0.02%. For valid comparison, PETase and STAR reactions were incubated at their respective optimal temperatures of 30°C and 40°C. After 24 hours incubation, STAR PETase showed ~ 1.6 and 3.9-fold improved yield of hydrolysis products (TPA+MHET+BHET) over PETase for respective degradation of powder and film substrates (Fig. 2A, B). Despite the slight increase in T_m_, the K95A mutation notably reduced activity in PETase by ~3.5-fold using PET powder substrate after 24 hours as previously described^31^(Fig. 2A). Conversely, in the STAR background, the K95A mutation increased activity by ~1.3 fold. Whilst activity in both backgrounds was unaffected by the R132N mutation, the K95A/R132N double mutation reduced PETase activity by ~13 fold. However, activity of STAR further increased by ~1.5-fold with this double mutation to yield the most active variant (~2.2-fold increased products yield over PETase). Interestingly, at the 72-hour time point the deleterious effects of the K95A and K95A/R132N mutations were notably reduced in PETase background. At 72 hours, activities of all STAR mutants exceeded STAR, with exception of STAR-R132N which displayed similar activity. The best performing enzyme, STAR^K95A/R132N^ degraded 100% of the powder substrate compared to 52% for PETase. For lcPET disk substrate, essentially the same phenotype was seen for mutants in the PETase background at 24 hours. Notably, whilst STAR^K95A^ showed improved activity on powdered substrate, its activity was significantly reduced on film substrate (~1.4 fold) compared to STAR (Fig. 2B). At 72 hours, the same trend was seen for most mutants in PETase/STAR backgrounds. STAR^K95A/R132N^ however shows significantly improved degradation over STAR at this later time point (~2 fold), degrading 7.2 % of the film substrate compared to 1.1 % for PETase. Surface morphology reflected the increased hydrolysis, with inter-connected depressions flanked by prominent ridge-like features seen for STAR^K95A/R132N^ that are absent for PETase, where discrete small crater-like features are prominent (Fig. 3). We also introduced the R280A mutation, shown to extend a substrate binding subsite and increase PETase hydrolysis by up to 32% ^14^, into the STAR and STAR^K95A/R132N^ backgrounds. Whilst this mutation increased activity of STAR on both substrates after 72 hours, no significant effect was observed in the STAR^K95A/R132N^ background.

**Figure 2.**
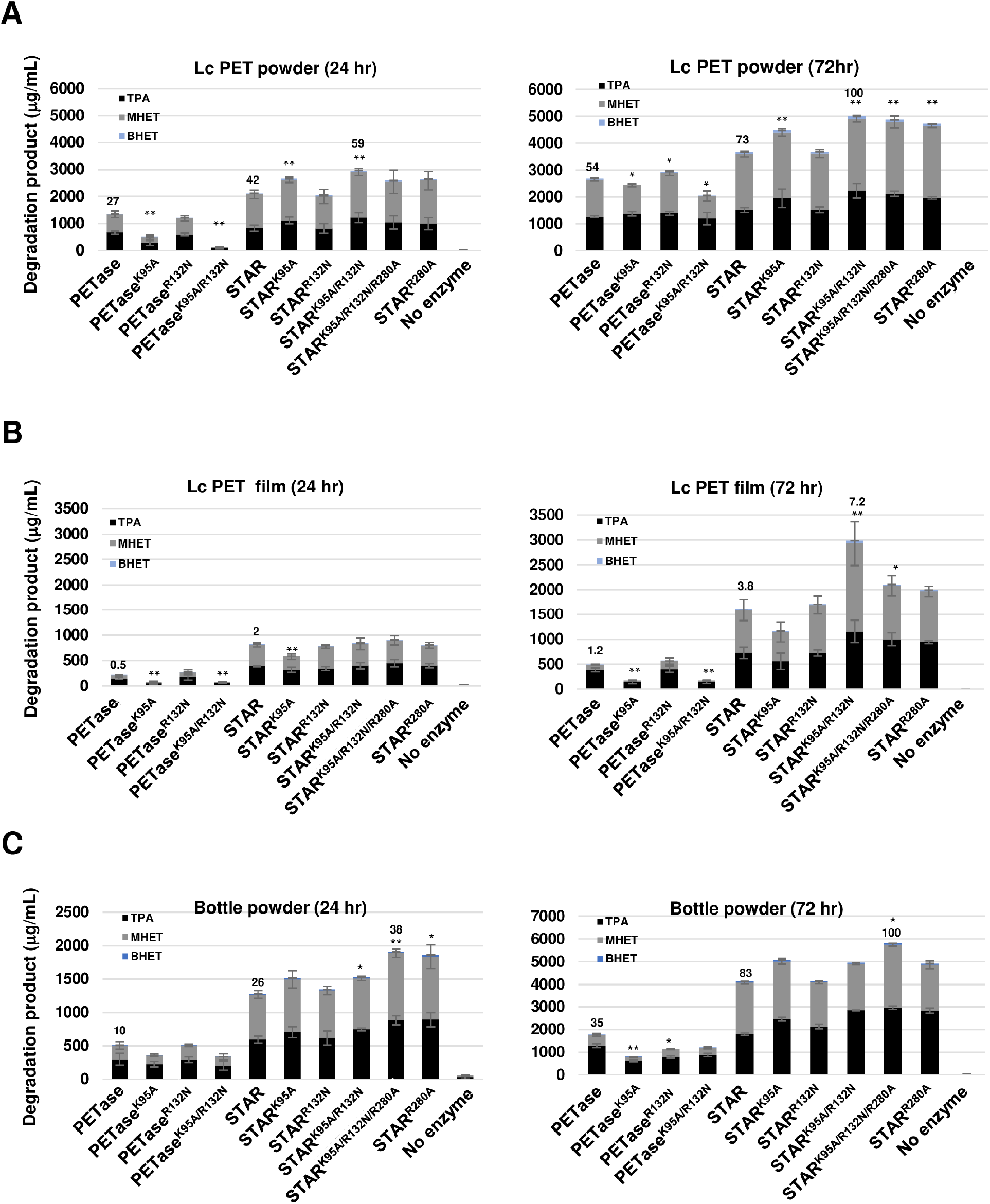
PET hydrolysis products generated by indicated enzymes after 24 and 72 hours incubation. PETase and STAR reactions were respectively incubated at 30°C and 40°C. n=3-5 ± SD. (A) LcPET powder. (B) LcPET film. (C) Commercial grade bottle powder. \**P*<0.05, \*\**P*<0.01 (one-way ANOVA with Tukey’s multiple comparison test). TPA: Terephthalic acid, MHET: Mono(2-hydroxyethyl) terephthalic acid, BHET: bis(2-hydroxyethyl) terephthalate. Numbers above bars indicate percentage degradation of substrate.

**Figure 3.**
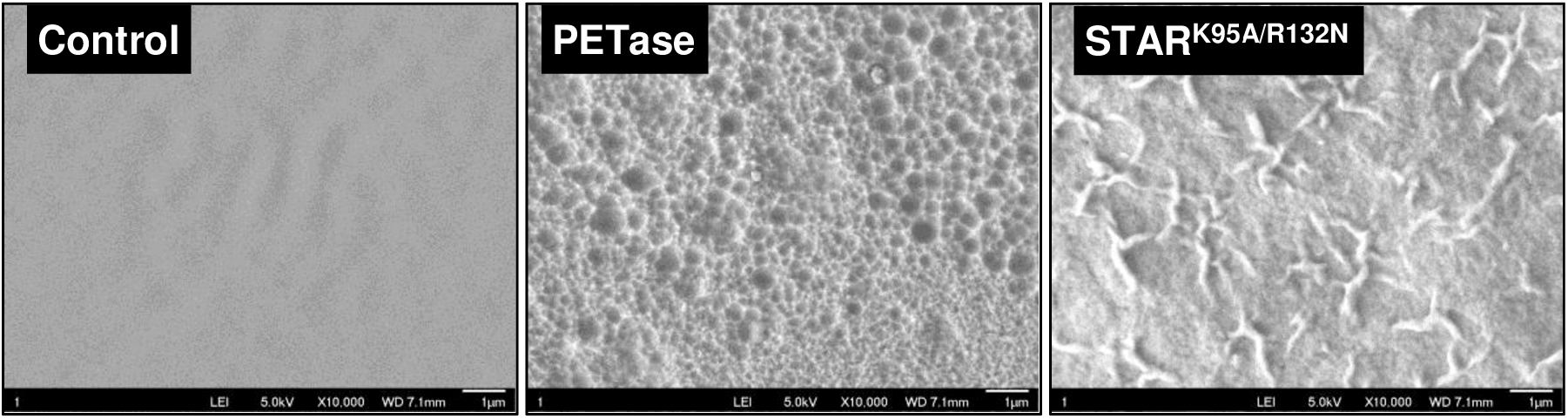
SEM images of enzymatically treated lcPET film substrate (24 hr).

We further assayed degradation of a micronized commercial grade PET bottle (9% crystallinity). PETase showed ~32% degradation at 72 hours, with the K95A mutations negatively impacting activity as before (Fig. 2C). STAR and STAR^R132N^ showed 83% degradation, whilst the best performing STAR^K95A/R132N/R280A^ variant (also termed STAR 4) degraded 100% substrate at this time point, 3-fold more than PETase. The other STAR variants also outperformed STAR, showing >90% degradation.

None of the variants investigated showed a different TPA to MHET product ratio to their parental scaffolds (Fig. S3). For the PETase panel, TPA contributed between 47 - 87% of total products. Whilst the STAR panel was more active overall, the TPA contribution was generally reduced to between 37-52% of total products. Given the inhibitory effect of accumulated MHET on PETases^39^, further engineering and/or supplementation with MHETase^40-42^ to convert MHET to TPA is warranted to yield higher amounts of the desired end product.

### Time course measurements show significant differences in hydrolysis rates

Reactions were next analysed at shorter time points using absorbance to measure PET powder hydrolysis^43,44^. Specific activity (SA) measured over the initial 4 hours for STAR reactions was 28-fold increased over PETase (Fig. 4A). The parental STAR enzyme clearly outperformed the STAR variants tested (0-4 hours), showing a specific activity (SA) of 229 μmol_TPAeq_ h^−1^ mg_enzyme-1_. After this lag phase, activities of all variants increased to yield similar amounts of degradation products to STAR after 8 hours. Introduction of the R280A mutation into STAR particularly curtailed PET hydrolysis over the initial 4 hours (45 μmol_TPAeq_ h^−1^ mg_enzyme_^−1^). Interestingly, this kinetic lag was most pronounced for enzyme variants comprising the R280A mutation (STAR^K95A/R132N/R280A^, STAR^R280A^). The K95A mutation notably reduced SA by up to 95% over 8 hours in the PETase backgrounds, whilst R132N inhibited activity by ~80%. Both mutations showed minimal perturbation of product yield in the STAR background over the same time period. At later time points (24-72 h), activity measurements followed the trend observed using HPLC to measure degradation (Fig 4B). With exception of the R132N point mutant, all other STAR variants out-performed STAR.

**Figure 4.**
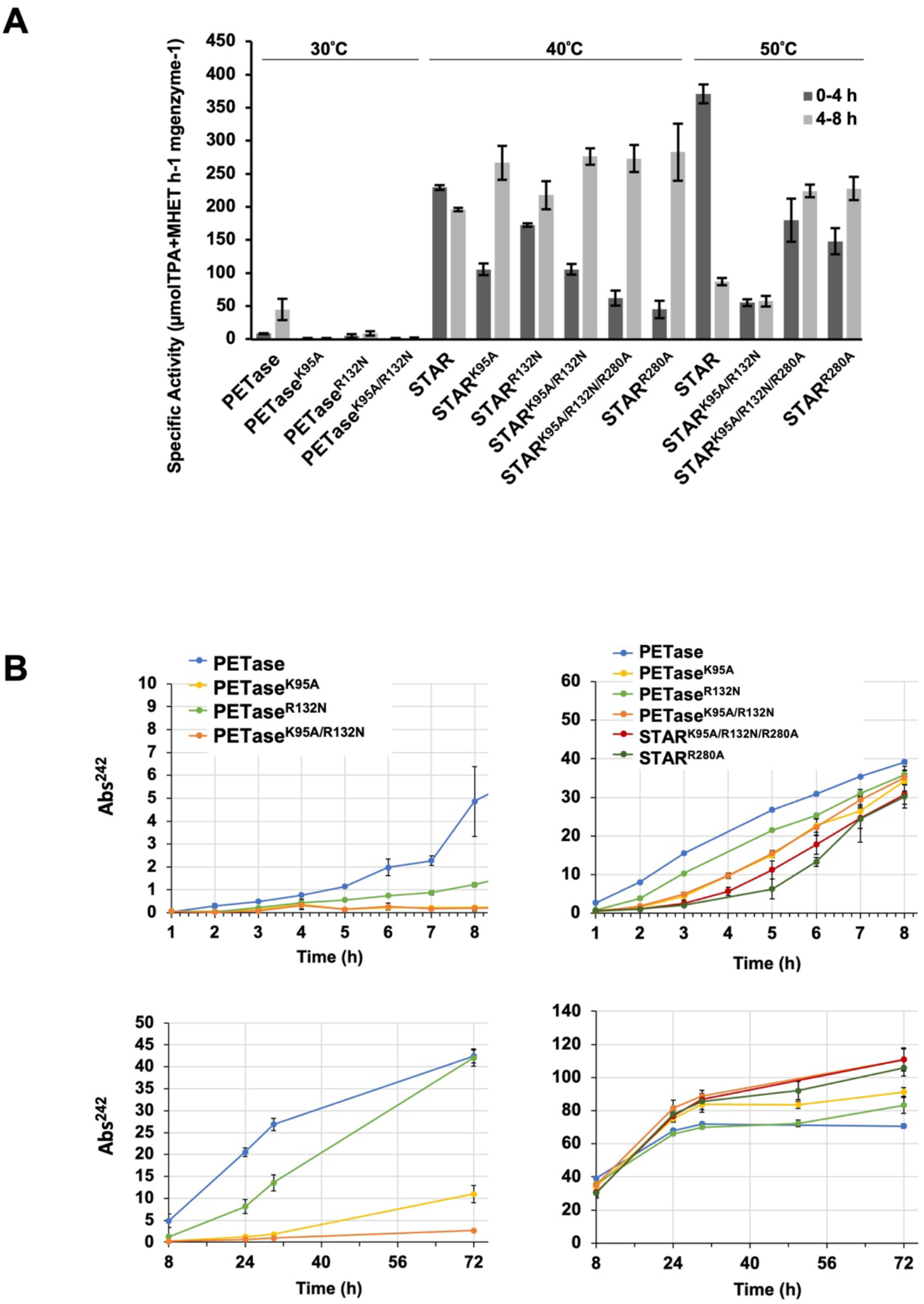
Specific activity measurements. (A) Specific activities of indicated enzymes measured using lcPET powder substrate. Reactions were incubated at indicated temperatures. (B) Time course measurement of lcPET powder hydrolysis over 8 hours. n=3 ± SD.

At a higher reaction temperature of 50°C, the SA of STAR increased 1.8-fold to 371 μmol_TPAeq_ h^−1^ mg_enzyme_^−1^, with similar increases observed for the STAR^K95A/R132N/R280A^ and STAR^R280A^ variants (measured over first four hours of reaction) (Fig. 4A, 5). However, activity tapered off between 3-8 hours, likely due to protein denaturation at the higher temperature closer to their respective T_m_s (57 - 62°C) (Table 1). Hence the 40°C optimal temperature of these enzymes for longer term reactions (>24 hours). Interestingly, despite a similar T_m_, STAR^K95A/R132N^ was very clearly less active at 50°C compared to 40°C over 8 hours reaction time (Fig. 5). This phenotype was reversed by introduction of the R280A mutation.

**Figure 5.**
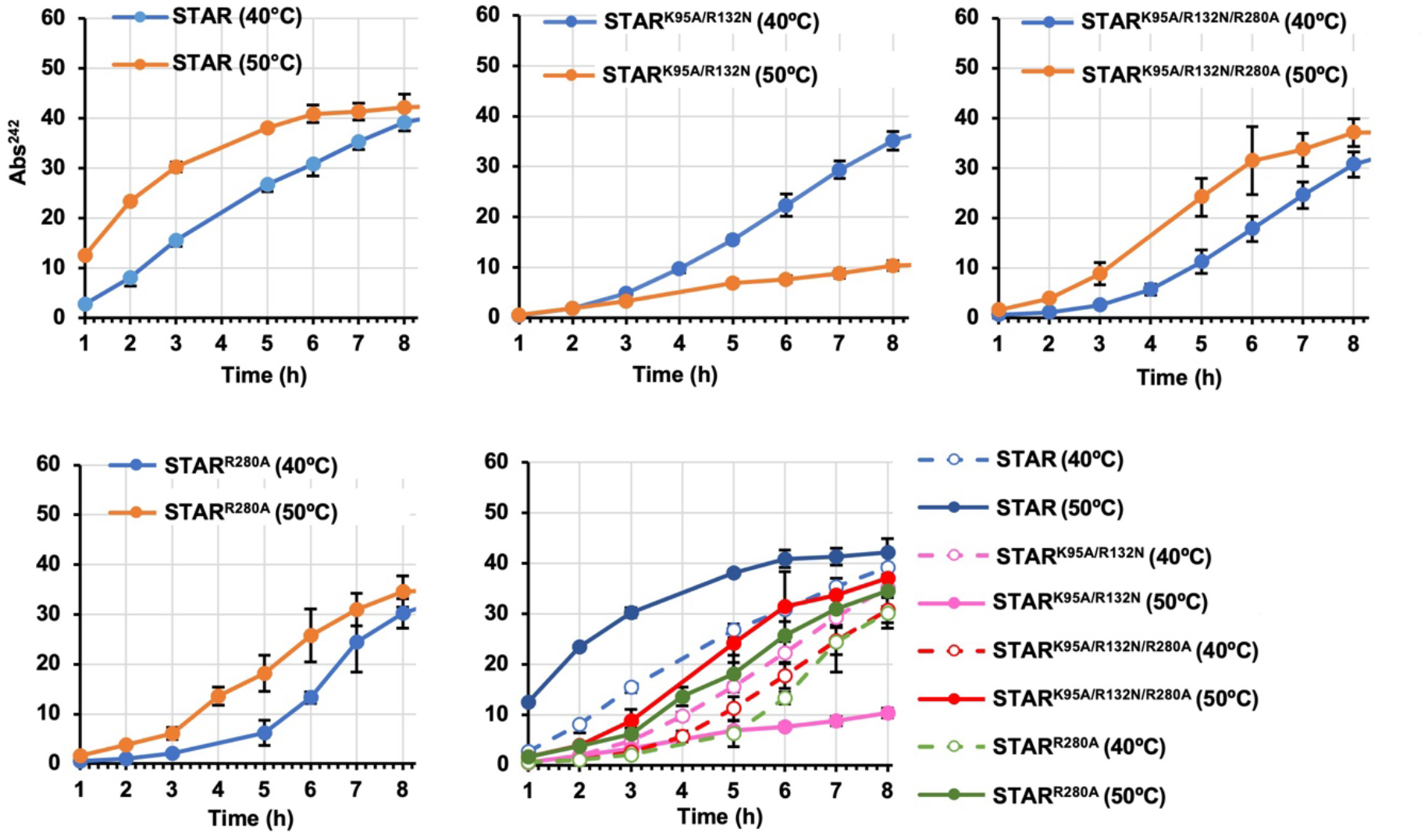
Time course measurement of lcPET powder hydrolysis over 8 hours for indicated enzymes at 40°C and 50°C. n=3 ± SD.

### Investigating surface charge and concentration-dependent PETase inhibition

PETase displays a strong concentration-dependent inhibition on PET film substrates, likely due to crowding conditions that promote non-productive inter-enzyme interactions^24,45^. Crystal forms of PETase have been observed wherein the R132 sidechain acts as a pseudo-substrate, docking into the substrate binding groove close to the catalytic triad of an adjacent protomer^46,47^. We therefore questioned whether R132 contributes to the inhibitory effect of high enzyme concentrations on catalytic activity. Inhibition was seen at enzyme concentrations >200 nM for both PETase and STAR on film substrate (Fig. 6). At higher concentrations (600 – 800 nM) the R132N mutation resulted in increased levels of inhibition in both PETase and STAR (relative to activity at 200 nM). This effect was more pronounced in STAR^R132N^, particularly at 800 nM where activity was reduced 25-fold compared to 8-fold in PETase^R132N^. It is possible that asparagine docks more favourably than lysine into the PETase and STAR active site regions. The S238 and E119 residues mutated in STAR are adjacent to the binding groove which the R132 sidechain occupies in crystal structures (Fig. S4). These mutations may afford a conformation that further favours asparagine insertion, contributing to enhanced inter-molecular binding and stronger inhibition.

**Figure 6.**
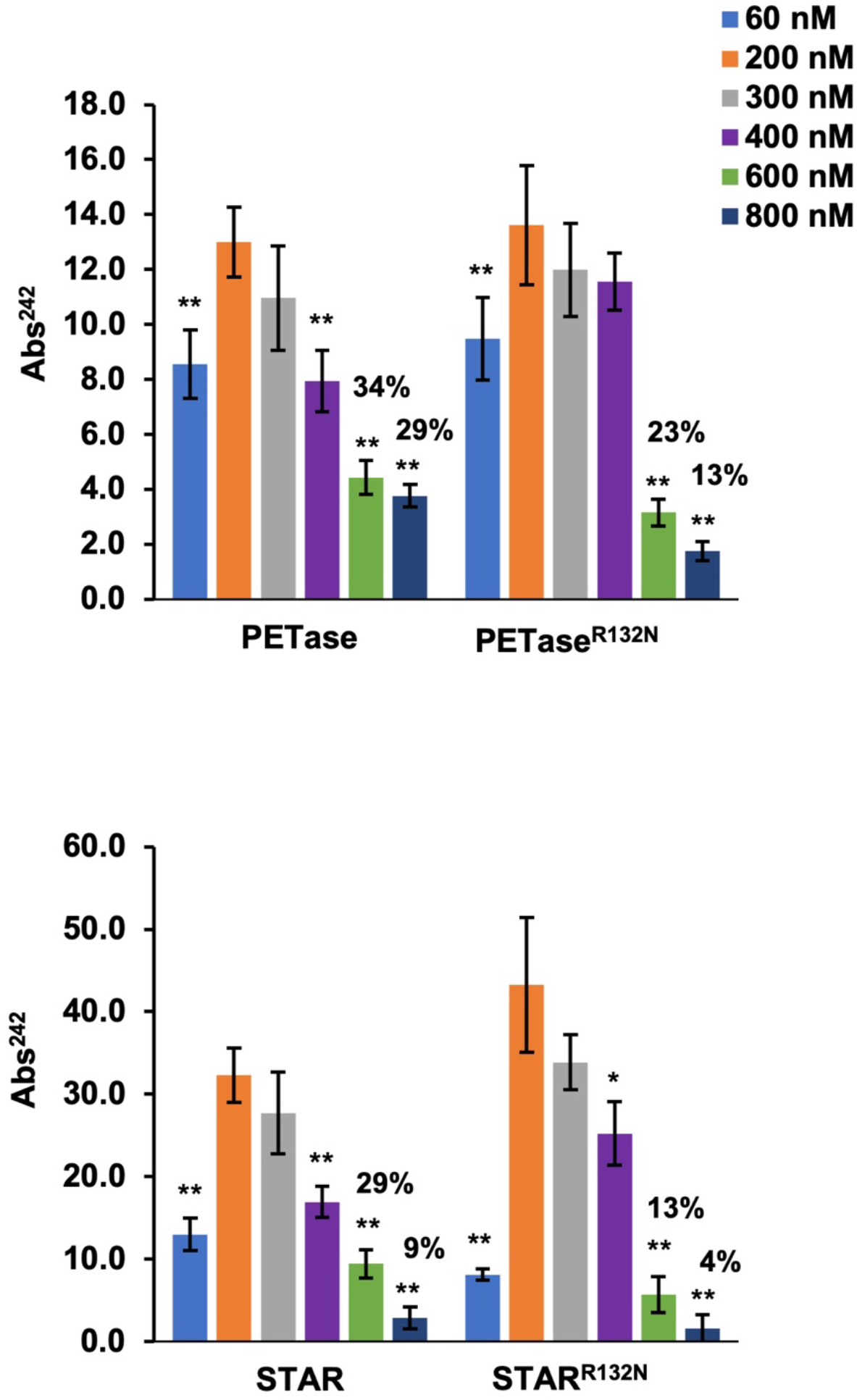
LcPET film hydrolysis over 24 hours for indicated enzymes. PETase and STAR reactions were respectively incubated at 30°C and 40°C. Percentage values indicate activity relative to 200 nM enzyme concentration. n=3 ± SD. \**P*<0.05, \*\**P*<0.005 (one-way ANOVA with Tukey’s multiple comparison test).

### Molecular dynamics simulations

The structural model of STAR-PETase shows introduction of a disulphide bridge connecting C282 and C233 (Fig. 7), further stabilizing the loop presenting the catalytic H127 and likely accounting for the increased thermostability measured (Table 1). The solvent exposed K95 resides at the inflection of a kinked α helix, the upper part of which contributes to the lining of the substrate binding channel in the vicinity of the active site (Fig. 7B). Mutation to alanine is predicted to both stabilize the α helix and potentially introduces stabilizing contacts with S242 in PETase and S242/W238 in STAR PETase, accounting for the increased T_m_ observed (Fig. 7B, Fig. S5A) (Table 1). This would rigidify the catalytic site (Fig. 7C, Fig. S5B), potentially restricting sampling of the multiple substrate conformations presented by a PET film substrate (mobile/rigid amorphous and crystalline regions). Hence the reduced activity on PET film measured for both STAR and PETase when the K95A mutation is introduced (Fig. 2). The W238 in STAR could form additional stacking interactions with the terephthalate ring (Fig. 7A), potentially increasing substrate affinity. Pre-stabilisation of its ring through A95-mediated contacts could favor more productive binding to PET powder substrate and increased activity as measured. R132 is presented on an extended α helix on the opposite face of the enzyme to the active site. Its side chain makes contacts with S61 from an adjacent loop and S136 on the same α helix. The S61 interaction is lost upon mutation to asparagine (Fig. 7B), increasing flexibility of the loop and potentially reducing T_m_ as observed (TABLE 1). Simulations predict this flexibility to propagate to the active site region considerably more in the STAR background compared to PETase (Fig. S5B), which could also contribute to the enhanced concentration-dependent inhibition observed in STAR-R132N (Fig. 6). The net effect of K95A and R132N combined is an active site with flexibility similar to that of STAR (Fig 6C). This rescue of active site flexibility could account for the ~3-fold improved digestion of PET film by STAR^K95/R132N^ over STAR^K95N^ (Fig. 2B). Kinetics experiments using powder substrate showed significantly reduced activity of STAR^K95A/R132N^ at 50°C compared to 40°C, in contrast to STAR (Fig 4). This phenotype was reversed by addition of the thermostabilising R280A mutation, again suggesting increased active site flexibility in STAR^K95A/R132N^ that does not favour catalysis at higher temperatures^48,49^.

**Figure 7.**
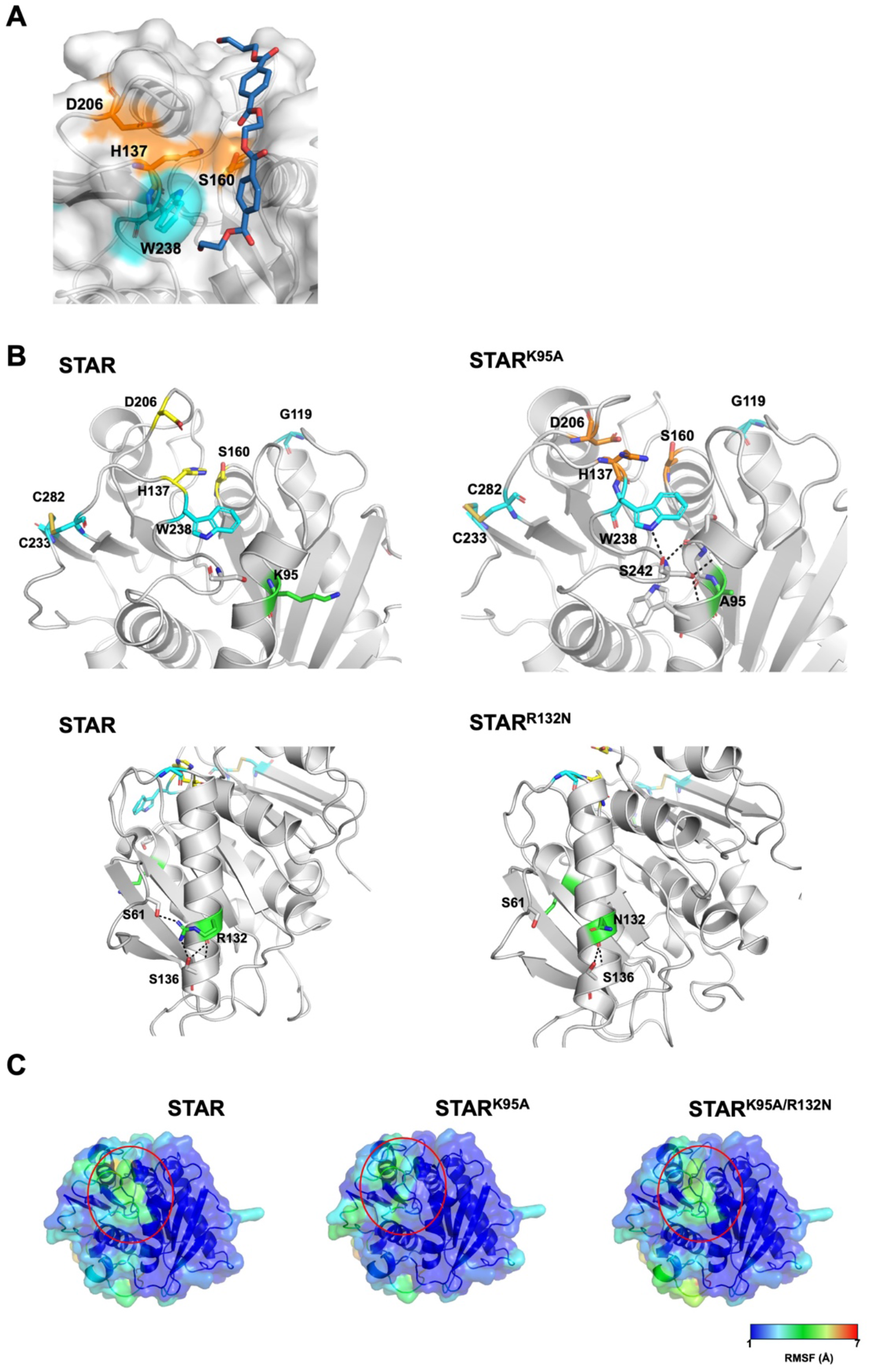
Molecular dynamics (MD) simulations predict modulation of active site flexibility by K95A and R132N mutations. (A) Proximity of W238 indole ring in STAR to PET terephthalate ring favours stacking interactions. Catalytic triad residues shown in orange. (B) The K95A mutation in STAR yields additional contacts with nearby residues S242 and W238. The R132N mutation results in loss of contact with S61. Catalytic triad residues depicted in yellow. Residues mutated in STAR are shown blue. (C) Flexibility of circled active site region (root mean square fluctuations) is reduced in STAR by addition of K95A and can be restored by further installation of R132N mutation.

## Discussion

In the PETase enzyme 18% of surface residues are charged, with ~4-fold increased cationic residues over anionic. These predominate on the enzyme face engaging with substrate (Fig. 1), and likely represent an evolutionary adaption towards digestion of recalcitrant PET substrate. As shown, removal of positive charge at the K95 position proximal to the active site, significantly reduced enzyme activity on all PET substrates. Substitution of the corresponding glutamic acid residue in a metagenome-derived thermostable cutinase (PET2) to lysine both increased T_m_ and amorphous PET hydrolysis activity by 1.4-fold ^50,51^, further validating a functional role of positive charge at this position. Recently, sequence reconstruction yielded a panel of thermostable, yet less active ancestral PETase variants, all of which were mutated at K95 to remove charge^26^. Notably, the same mutation in context of the thermostabilized STAR variant increased activity on PET powder yet reduced activity on PET film, reiterating the need to engineer and select enzymes using a substrate type/morphology similar or identical to anticipated commercial feedstock^52^. MD simulations suggest the context-dependent phenotype is a result of an additional enzyme-substrate interaction mediated by the S238W mutation in STAR that can be further enhanced by the K95A mutation. We speculate that tryptophan side chain mobility is constrained to both reduce entropic cost of substrate binding and promote interaction with the predominantly mobile amorphous regions in mechanically processed and low crystallinity PET substrate. Active site flexibility likely facilitates broader conformational selectivity towards rigid PET chains^30^. Hence whilst the additional rigidity in STAR^K95A^ does not aid in hydrolysis of higher crystallinity PET film substrates (Fig. 2B), it can be relieved by further installation of the R132N mutation which is predicted to allosterically increase active site flexibility. Notably, the S238A mutation is reported to change the active site’s conformational landscape and alter substrate selectivity^53^, indicating a role for the S238 region in modulating enzyme dynamics. The R132E mutation has also been shown to increase activity in the context of the thermostabilised Is-8p variant^25^, further highlighting effects of charge modification at this distal residue. The R132N mutation also potentiated the characteristic concentration-dependent inhibition of enzyme activity on PET film, particularly in STAR PETase. This likely arises from the increased enzyme flexibility that favours non-productive inter-molecular interactions at high enzyme concentrations. One of these interactions is possibly recapitulated in crystal structures, where the R132 side chain of one monomer is seen inserting into the active site region of another. Further structural studies are required to delineate precise mechanisms of action.

In summary, we have characterized the thermostable STAR PETase variant with improved activity over the parental enzyme and further improved its performance through surface charge engineering. The effects of surface charge mutation are shown to be dependent both on enzyme structural context and PET substrate morphology. The STAR4 variant completely degraded a pre-processed commercial bottle substrate in 3 days at 40°C. Whilst the comparatively low temperature optimum of STAR4 may contribute to reduced conversion costs at scale, its specific activity is 2-4 fold lower than other PETase mutants described^44^. As depolymerization rate is a key determinant of industrial scalability^44^ further enzyme engineering of STAR4 will likely be required.

## Methods

### Cloning

The *Is*PETase gene was codon-optimized for recombinant expression in *Escherichia coli* and synthesized (IDT). The gene was amplified using primers 1a and 1b (see Supporting Information: Table S1 for list of oligonucleotide primers). The purified PCR product was cloned with a C-terminus 6-His tag into pET22b vector via NdeI/XhoI using the In-Fusion HD Cloning kit following the manufacturer’s protocol (Takara Bio). The construct was transformed into chemically competent *E. coli* DH5α cells (Invitrogen) and the correct sequence was confirmed by sequencing.

The variant constructs were made by introducing K95A, R132N and R280A point mutations in the PETase and STAR PETase constructs by PCR, using primers 2a and 2b, 3a and 3b, 4a and 4b. The PCR products were digested using DpnI enzyme to remove parental templates and transformed into chemically competent *E. coli* DH5α cells (Invitrogen) and the correct sequence was confirmed by sequencing.

### Recombinant protein expression and purification

Plasmid constructs of PETase/ variants and STAR variants were transformed into *E. coli*. Shuffle T7 competent cell following the manufacturer’s Protocol (New England Biolabs) and grown on LB/agar plates containing 0.1 mg/mL ampicillin at 30°C for 2 days. Overnight cultures were prepared by inoculating single colonies in LB broth containing 0.1 mg/mL ampicillin and grown at 30°C with constant shaking. For over-expression, the overnight cultures were diluted 80-fold in the same media, grown at 30°C until OD600=0.5, and induced using 0.5mM IPTG. The culture was grown at 16°C for 20 h. The proteins were purified using Ni-NTA column (Cytiva) following manufacturer’s protocol. The purified proteins were concentrated and buffer exchanged into PBS using 10 kDa MWCO Ultra centrifugal filters (Amicon). The concentration and purity of the purified proteins were determined by OD280 using Nanodrop and SDS/PAGE respectively.

### PET depolymerization assays

Degradation of PET studies were carried out using low crystallinity lcPET powder, lcPET film and pretreated pc-PET bottle powder as substrate. The lcPET powder was prepared by roughly cutting PET 1445 (Goodfellow™) into cm-sized coupons. About 15-20 grams of coupons were loaded in the aluminium bowl of A11 basic analytical mill (IKA™) and allowed to rest in liquid nitrogen for 15 seconds followed by grinding for 30 seconds. This step of liquid nitrogen addition and grinding was repeated five times. The ground powder was sieved using steel wire mesh (WS Tyler™) no. 140 (106 microns) and particles smaller than 106 microns were used for the degradation experiments. The crystallinity of these particles is around 8% as measured by DSC. LcPET 1445 film was hole-punched to give circular disks (6mm in diameter, 0.28 square centimetre and 8.5 ± 0.2 mg/disk). The crystallinity of the film is around 7% as measured by DSC. pc-PET bottle powder was prepared from clean pc-PET water bottles with caps and labels removed and roughly cut into cm-sized coupons. Approximately 2 grams were weighed and sealed flat in aluminium foil pocket. Each pocket was heated using hot plate (260°C) and pressed with a weight of 500g for 1 minute followed by heating for 5 minutes. The samples were quenched in ice bath for 5 minutes. Sample was taken out of the aluminium pocket and washed with DI water and dried with a wipe. Heat-treated PET (15-20 grams) was loaded in the aluminium bowl of A11 basic analytical mill (IKA™) and ground for 30 seconds and rested for 1 minutes and this cycle was repeated 5 times. The ground powder was sieved using steel wire mesh to obtain particles with 150-300 microns size distribution which were used for enzymatic degradation. The crystallinity of this powder is around 9% as measured by DSC. Enzyme reactions (200 μl) were carried out using either 1 mg substrate powder or 8.5 ± 0.2 mg disk, 200 nM enzyme in 100 mM sodium phosphate buffer pH 8.0. Reactions were performed for 1 and 3 days. The enzymes were inactivated by heating at 95°C for 10 minutes before centrifuged at 13,000 rpm for 10 minutes. Supernatants were analysed by HPLC to quantify TPA, MHET and BHET products. Triplicate reactions were setup for each enzyme/ variant.

### PET depolymerization analysis by ultraviolet (UV) light absorbance

Absorbance readings at 242 nm were used to detect released TPA, MHET and BHET upon PET depolymerization following established protocols^43,54^. 1.5 μL reaction samples were analysed using Nano Drop Eight Spectrophotometer (Thermoscientific). No-enzyme control reaction was used for instrument blanking and “automated pathlength function” was selected to record absorbance values based on pathlength of 1 cm. Concentration of PET degradation products was calculated using average extinction coefficient of TPA and MHET (13,373 M^−1^ cm^−1^) following Beer-Lambert law. Individual TPA (11,762 M^−1^ cm^−1^) and MHET (14,985 M^−1^ cm^−1^) extinction coefficients were determined from standard curves generated with compounds diluted in 100 mM (pH 8.0) sodium phosphate buffer in the range of 0 μM to 20,000 μM. BHET was excluded from calculations due to significantly lower levels observed in PET degradation reactions. Samples with absorbance readings exceeding instrument detection range were diluted in the same 100 mM sodium phosphate buffer.

### Kinetics Assay

Kinetics assays were performed to determine specific activity of all enzyme variants following established protocols^43,54^. lcPET powder degradation reactions (1 mL) were carried out in 1.5 mL Eppendorf tubes at optimum temperatures of corresponding enzymes, unless stated otherwise. Final enzyme concentration in all reactions was set to 60 nM. Enzyme stocks (3 μM) in 20 μL volumes were added to 2 mg lcPET powder pre-heated in 980 μL sodium phosphate buffer (100mM, pH 8.0). PET depolymerization was assessed using UV light absorbance at 242 nm as described. Samples were taken every hour from 0-8 h and additional readings were taken at 24 h, 30 h and 72 h. Specific activity of enzymes was determined from accumulated PET degradation products in 0-4 h and 4-8 h time periods.

### Differential scanning fluorimetry assay

Protein samples were diluted in PBS to 5 μM and mixed with 5000× SYPRO Orange Protein Gel Stain (Invitrogen) to final 1.25x concentration. The mixtures (50 μL) were incubated at RT for 20 mins and assayed using HEX channel on BioRad CFX96 system. Fluorescence readings were taken at 0.5°C sec^−1^ increments from 25°C to 95°C.

### Molecular dynamics simulations

The crystal structure of *I. sakaiensis* PETase (PDB ID 5XJH; resolution 1.5 Å^14^) (referred to as PETase) was used to model the structure of PET^K95A^, PET^R132N^, PET^K95A/R132N^. The STAR enzyme has 4 point mutations (G119Q, N233C, S238W and S282C) as compared to the PETase. The crystal structure of PETase was modified to construct the STAR by mutating the 4 corresponding residues. A di-sulphide bridge between the N233C and S282C was introduced guided by the observation that the homologous positions in the crystal structure (PDB ID 8H5M; resolution 1.9 Å) of *P. sakaiensis* PETase are engaged in a disulphide. STAR was used to generate the STAR^K95A^, STAR^R132N^ and STAR^K95A/R132N^ by mutating the corresponding residues.

The 3D structure of the linear PET molecule was built using the *Maestro* module and minimized using the *Macromodel* module employing the OPLS-2005 force field^55^ in the program Schrodinger 12.0^56^. The PET molecule was then prepared with *Ligprep* module in Schrodinger that generates low energy tautomers and enumerates realistic protonation states at physiological pH. The prepared PET was then docked into the binding pocket on the surface of PET^WT^ and STAR^WT^ using the *Glide*^57^ module of Schrodinger. A box of size 10 x 10 x 10 ? for molecular docking centred on the selected active site residues (S131, H208, D177) was used to confine the search space of PET molecule. For the grid generation, the default *Glide* settings were used. Docking was carried out using standard protocols^58^ and the docked conformation of each ligand was evaluated using the *Glide* Extra Precision (XP) scoring function.

The modelled PETase and STAR and their mutant versions together with the PETase – PET and STAR-PET complexes were subject to Molecular Dynamics (MD) simulations. MD simulations were carried out with the *pmemd*.*cuda* module of the program Amber18^59^. The *Antechamber* module in Amber was used to generate the partial charges and force field parameters for the PET molecule. All atom versions of the Amber14SB force field (ff14SB) ^60^ and the general Amber force field (GAFF) ^61^ were used to model the protein and the PET molecule respectively. The *Xleap* module of Amber was used to prepare the systems for the MD simulations. All the simulation systems were neutralized with appropriate numbers of counter ions. Each neutralized system was solvated in an octahedral box with TIP3P^62^ water molecules, leaving at least 10 Å between the solute atoms and the borders of the box. MD simulations were all carried out in explicit solvent at 300 K. During the simulations, the long-range electrostatic interactions were treated with the particle mesh Ewald^63^ method using a real space cut off distance of 9Å. The Settle^64^ algorithm was used to constrain bond vibrations involving hydrogen atoms, which allowed a time step of 2fs during the simulations. Solvent molecules and counter ions were initially relaxed using energy minimization with restraints on the nonhydrogen atoms of the enzyme-PET complex structures. Unrestrained energy minimization was next applied to remove any steric clashes. The system was next gradually heated from 0 to 300 K using MD simulations with positional restraints (force constant: 50 kcal mol^−1^ Å^−2^) on the nonhydrogen atoms of the enzyme-PET complex structures over a period of 0.25 ns allowing water molecules and ions to move freely. For the duration of an additional 0.25 ns, the positional restraints were gradually reduced followed by a 2 ns unrestrained MD simulation to equilibrate all the atoms. Production MD simulations were finally carried out on each system for 100 ns in triplicates. Accelerated MD (aMD) simulations was used to enhance the conformational sampling of the PETases and STAR enzymes during the MD simulations^65^. aMD runs (dual boost potentila) for 250 ns were carried in triplicates. Simulation trajectories were visualized using VMD^66^ and figures were generated using PyMOL^67^.

## Author Contributions

D.K., Z.L., B.S., C.V., S.L., J.G designed the project and experiments. D.K., Z.L., B.S., V.V.S., A.A.,

R.R. carried out experiments. S.R. and C.V. carried out molecular dynamics simulations. S.L. secured funding for the project. J.G. wrote manuscript.

## Data availability

The datasets generated during and/or analysed during the current study are available from the corresponding author on reasonable request.

## References

1 Rolf-Joachim Müller, H. S., Jörn Profe, Karolin Dresler, Wolf-Dieter Deckwer. Enzymatic Degradation of Poly(ethylene terephthalate): Rapid Hydrolyse using a Hydrolase from T. fusca. Macromolecular Rapid Communications 26, 1400–1405 (2005).

2 Yoshida, S. et al. A bacterium that degrades and assimilates poly(ethylene terephthalate). Science 351, 1196 (2016). 10.1126/science.aad6359

3 Oda, K. & Wlodawer, A. Development of Enzyme-Based Approaches for Recycling PET on an Industrial Scale. Biochemistry (2024). 10.1021/acs.biochem.3c00554

4 Geyer, R., Jambeck, J. R. & Law, K. L. Production, use, and fate of all plastics ever made. Sci Adv 3, e1700782 (2017). 10.1126/sciadv.1700782

5 Shi, L. et al. Complete Depolymerization of PET Wastes by an Evolved PET Hydrolase from Directed Evolution. Angew Chem Int Ed Engl 62, e202218390 (2023). 10.1002/anie.202218390

6 Son, H. F. et al. Rational Protein Engineering of Thermo-Stable PETase from Ideonella sakaiensis for Highly Efficient PET Degradation. ACS Catalysis 9, 3519–3526 (2019). 10.1021/acscatal.9b00568

7 Pasula, R. R., Lim, S., Ghadessy, F. J. & Sana, B. The influences of substrates’ physical properties on enzymatic PET hydrolysis: Implications for PET hydrolase engineering. Eng Biol 6, 17–22 (2022). 10.1049/enb2.12018

8 Cui, Y. et al. Computational Redesign of a PETase for Plastic Biodegradation under Ambient Condition by the GRAPE Strategy. ACS Catalysis 11, 1340–1350 (2021). 10.1021/acscatal.0c05126

9 Lu, H. et al. Machine learning-aided engineering of hydrolases for PET depolymerization. Nature 604, 662–667 (2022). 10.1038/s41586-022-04599-z

10 Tournier, V. et al. An engineered PET depolymerase to break down and recycle plastic bottles. Nature 580, 216–219 (2020). 10.1038/s41586-020-2149-4

11 Thiyagarajan, S., Maaskant-Reilink, E., Ewing, T. A., Julsing, M. K. & van Haveren, J. Back-to-monomer recycling of polycondensation polymers: opportunities for chemicals and enzymes. RSC Advances 12, 947–970 (2022). 10.1039/D1RA08217E

12 Austin, H. P. et al. Characterization and engineering of a plastic-degrading aromatic polyesterase. Proceedings of the National Academy of Sciences 115, E4350 (2018). 10.1073/pnas.1718804115

13 Bell, E. L., Smithson, R., Kilbride, S. et al. Directed evolution of an efficient and thermostable PET depolymerase. Nature Catalysis 5, 673–681 (2022). 10.1038/s41929-022-00821-3

14 Joo, S. et al. Structural insight into molecular mechanism of poly(ethylene terephthalate) degradation. Nature Communications 9, 382 (2018). 10.1038/s41467-018-02881-1

15 Liu, B. et al. Protein crystallography and site-direct mutagenesis analysis of the poly(ethylene terephthalate) hydrolase PETase from Ideonella sakaiensis. Chembiochem (2018). 10.1002/cbic.201800097

16 Jia, Y. et al. Hydrophobic cell surface display system of PETase as a sustainable biocatalyst for PET degradation. Front Microbiol 13, 1005480 (2022). 10.3389/fmicb.2022.1005480

17 Samak, N. A. et al. Recent advances in biocatalysts engineering for polyethylene terephthalate plastic waste green recycling. Environ Int 145, 106144 (2020). 10.1016/j.envint.2020.106144

18 Meng, X. et al. Protein engineering of stable IsPETase for PET plastic degradation by Premuse. Int J Biol Macromol 180, 667–676 (2021). 10.1016/j.ijbiomac.2021.03.058

19 Rennison, A., Winther, J. R. & Varrone, C. Rational Protein Engineering to Increase the Activity and Stability of IsPETase Using the PROSS Algorithm. Polymers (Basel) 13 (2021). 10.3390/polym13223884

20 Yin, Q., You, S., Zhang, J., Qi, W. & Su, R. Enhancement of the polyethylene terephthalate and mono-(2-hydroxyethyl) terephthalate degradation activity of Ideonella sakaiensis PETase by an electrostatic interaction-based strategy. Bioresour Technol 364, 128026 (2022). 10.1016/j.biortech.2022.128026

21 Zurier, H. S. & Goddard, J. M. A high-throughput expression and screening platform for applications-driven PETase engineering. Biotechnol Bioeng 120, 1000–1014 (2023). 10.1002/bit.28319

22 Sana, B. et al. Thermostability enhancement of polyethylene terephthalate degrading PETase using self- and nonself-ligating protein scaffolding approaches. Biotechnol Bioeng 120, 3200–3209 (2023). 10.1002/bit.28523

23 Elizabeth L. Bell, R. S., Siobhan Kilbride, Jake Foster, Florence J. Hardy, Saranarayanan Ramachandran, Aleksander A. Tedstone, Sarah J. Haigh, Arthur A. Garforth, Philip J. R. Day, Colin Levy, Michael P. Shaver & Anthony P. Green Directed evolution of an efficient and thermostable PET depolymerase. Nature Catalysis 5, 673–681 (2022).

24 Avilan, L. et al. Concentration-Dependent Inhibition of Mesophilic PETases on Poly(ethylene terephthalate) Can Be Eliminated by Enzyme Engineering. ChemSusChem 16, e202202277 (2023). 10.1002/cssc.202202277

25 Lee, S. H. et al. Three-directional engineering of IsPETase with enhanced protein yield, activity, and durability. J Hazard Mater 459, 132297 (2023). 10.1016/j.jhazmat.2023.132297

26 Joho, Y. et al. Ancestral Sequence Reconstruction Identifies Structural Changes Underlying the Evolution of Ideonella sakaiensis PETase and Variants with Improved Stability and Activity. Biochemistry 62, 437–450 (2023). 10.1021/acs.biochem.2c00323

27 Erickson, E. et al. Sourcing thermotolerant poly(ethylene terephthalate) hydrolase scaffolds from natural diversity. Nat Commun 13, 7850 (2022). 10.1038/s41467-022-35237-x

28 Tiong, E. et al. Expression and engineering of PET-degrading enzymes from Microbispora, Nonomuraea, and Micromonospora. Applied and environmental microbiology 89, e0063223 (2023). 10.1128/aem.00632-23

29 Fecker, T. et al. Active Site Flexibility as a Hallmark for Efficient PET Degradation by I. sakaiensis PETase. Biophys J 114, 1302–1312 (2018). 10.1016/j.bpj.2018.02.005

30 Thomsen, T. B. et al. Rate Response of Poly(Ethylene Terephthalate)-Hydrolases to Substrate Crystallinity: Basis for Understanding the Lag Phase. ChemSusChem 16, e202300291 (2023). 10.1002/cssc.202300291

31 Furukawa, M., Kawakami, N., Oda, K. & Miyamoto, K. Acceleration of Enzymatic Degradation of Poly(ethylene terephthalate) by Surface Coating with Anionic Surfactants. ChemSusChem 11, 4018–4025 (2018). 10.1002/cssc.201802096

32 Sagong, H. Y. et al. Implications for the PET decomposition mechanism through similarity and dissimilarity between PETases from Rhizobacter gummiphilus and Ideonella sakaiensis. J Hazard Mater 416, 126075 (2021). 10.1016/j.jhazmat.2021.126075

33 Pedersen, J. N., Zhou, Y., Guo, Z. & Perez, B. Genetic and chemical approaches for surface charge engineering of enzymes and their applicability in biocatalysis: A review. Biotechnol Bioeng 116, 1795–1812 (2019). 10.1002/bit.26979

34 Tournier, V. et al. An engineered PET depolymerase to break down and recycle plastic bottles. Nature 580, 216–219 (2020). 10.1038/s41586-020-2149-4

35 Tournier, V. NOVEL ESTERASES and USES THEREOF. WO 2021/005199 (2021).

36 Austin, H. P. et al. Characterization and engineering of a plastic-degrading aromatic polyesterase. Proc Natl Acad Sci U S A 115, E4350–E4357 (2018). 10.1073/pnas.1718804115

37 Ye Zhou, B. P., Weiwei Hao, Jiaboa Lu, Renjun Gao, Zheng Guo. The additive mutational effects from surface charge engineering: A compromise between enzyme activity, thermostability and ionic liquid tolerance. Biochemical Engineering Journal 148, 195–204 (2019).

38 Matulis, D., Kranz, J. K., Salemme, F. R. & Todd, M. J. Thermodynamic stability of carbonic anhydrase: measurements of binding affinity and stoichiometry using ThermoFluor. Biochemistry 44, 5258–5266 (2005). 10.1021/bi048135v

39 Wei, R. et al. Engineered bacterial polyester hydrolases efficiently degrade polyethylene terephthalate due to relieved product inhibition. Biotechnol Bioeng 113, 1658–1665 (2016). 10.1002/bit.25941

40 Zhang, J. et al. Computational design of highly efficient thermostable MHET hydrolases and dual enzyme system for PET recycling. Commun Biol 6, 1135 (2023). 10.1038/s42003-023-05523-5

41 Barth, M. et al. A dual enzyme system composed of a polyester hydrolase and a carboxylesterase enhances the biocatalytic degradation of polyethylene terephthalate films. Biotechnol J 11, 1082–1087 (2016). 10.1002/biot.201600008

42 Knott, B. C. et al. Characterization and engineering of a two-enzyme system for plastics depolymerization. Proc Natl Acad Sci U S A 117, 25476–25485 (2020). 10.1073/pnas.2006753117

43 Zhong-Johnson, E. Z. L., Voigt, C. A. & Sinskey, A. J. An absorbance method for analysis of enzymatic degradation kinetics of poly(ethylene terephthalate) films. Scientific Reports 11, 928 (2021). 10.1038/s41598-020-79031-5

44 Arnal, G. et al. Assessment of Four Engineered PET Degrading Enzymes Considering Large-Scale Industrial Applications. ACS Catal 13, 13156–13166 (2023). 10.1021/acscatal.3c02922

45 Baath, J. A., Borch, K., Jensen, K., Brask, J. & Westh, P. Comparative Biochemistry of Four Polyester (PET) Hydrolases*. Chembiochem 22, 1627–1637 (2021). 10.1002/cbic.202000793

46 Han, X. et al. Structural insight into catalytic mechanism of PET hydrolase. Nature Communications 8, 2106 (2017). 10.1038/s41467-017-02255-z

47 Palm, G. J. et al. Structure of the plastic-degrading Ideonella sakaiensis MHETase bound to a substrate. Nat Commun 10, 1717 (2019). 10.1038/s41467-019-09326-3

48 Wang, W., Dasetty, S., Sarupria, S., Blenner, M. Rational engineering of low temperature activity in thermoalkalophilic Geobacillus thermocatenulatus lipase. Biochemical Engineering Journal 174, 108093 (2021).

49 Lam, S. Y., Yeung, R. C., Yu, T. H., Sze, K. H. & Wong, K. B. A rigidifying salt-bridge favors the activity of thermophilic enzyme at high temperatures at the expense of low-temperature activity. PLoS Biol 9, e1001027 (2011). 10.1371/journal.pbio.1001027

50 A. Nakamura, N. K., N. Koga and R. Lino. Positive Charge Introduction on the Surface of Thermostabilized PET Hydrolase Facilitates PET Binding and Degradation. ACS Catalysis 11, 8550–8564 (2021). 10.1021/acscatal.1c01204

51 Meilleur, C., Hupe, J. F., Juteau, P. & Shareck, F. Isolation and characterization of a new alkali-thermostable lipase cloned from a metagenomic library. J Ind Microbiol Biotechnol 36, 853–861 (2009). 10.1007/s10295-009-0562-7

52 Erickson, E. et al. Comparative Performance of PETase as a Function of Reaction Conditions, Substrate Properties, and Product Accumulation. ChemSusChem 15, e202102517 (2022). 10.1002/cssc.202102517

53 Boyang Guo, S. R. V., Ximena Lopez-Lorenzo, Patricia Saenz-Mendez, Sara Rönnblad Ericsson, Yuan Fang, Xinchen Ye, Karen Schriever, Eva Bäckström, Antonino Biundo, Roman A. Zubarev, István Furó, Minna Hakkarainen, and Per-Olof Syrén*. Conformational Selection in Biocatalytic Plastic Degradation by PETase. ACS Catal. 12, 3397–3409 (2022).

54 Arnal, G. et al. Assessment of Four Engineered PET Degrading Enzymes Considering Large-Scale Industrial Applications. ACS Catalysis 13, 13156–13166 (2023). 10.1021/acscatal.3c02922

55 Kaminski, G. A., Friesner, R. A., Tirado-Rives, J. & Jorgensen, W. L. Evaluation and reparametrization of the OPLS-AA force field for proteins via comparison with accurate quantum chemical calculations on peptides. Journal of Physical Chemistry B 105, 6474–6487 (2001). 10.1021/jp003919d

56 Schrodinger, version 9.0. Schrödinger LLC, New York, NY, 2009 (2009).

57 Friesner, R. A. et al. Glide: A new approach for rapid, accurate docking and scoring. 1. Method and assessment of docking accuracy. Journal of Medicinal Chemistry 47, 1739–1749 (2004). 10.1021/jm0306430

58 Kannan, S. et al. Small Molecules Targeting the Inactive Form of the Mnk1/2 Kinases. Acs Omega 2, 7881–7891 (2017). 10.1021/acsomega.7b01403

59 Case, D. e. a. Amber 18. University of California, San Francisco.

60 Maier, J. A. et al. ff14SB: Improving the Accuracy of Protein Side Chain and Backbone Parameters from ff99SB. Journal of Chemical Theory and Computation 11, 3696–3713 (2015). 10.1021/acs.jctc.5b00255

61 Wang, J. M., Wolf, R. M., Caldwell, J. W., Kollman, P. A. & Case, D. A. Development and testing of a general amber force field. Journal of Computational Chemistry 25, 1157–1174 (2004). 10.1002/jcc.20035

62 Jorgensen, W. L., Chandrasekhar, J., Madura, J. D., Impey, R. W. & Klein, M. L. Comparison of simple potential functions for simulating liquid water. Journal of Chemical Physics 79, 926–935 (1983). 10.1063/1.445869

63 Darden, T., York, D. & Pedersen, L. Particle mesh ewald - AN N.LOG(N) method for ewald sums in large systems. Journal of Chemical Physics 98, 10089–10092 (1993). 10.1063/1.464397

64 Miyamoto, S. & Kollman, P. A. Settle - An analytical version of the SHAKE and RATTLE algorithm for rigid water models. Journal of Computational Chemistry 13, 952–962 (1992). 10.1002/jcc.540130805

65 Hamelberg, D., Mongan, J. & McCammon, J. A. Accelerated molecular dynamics: a promising and efficient simulation method for biomolecules. J Chem Phys 120, 11919–11929 (2004). 10.1063/1.1755656

66 Humphrey, W., Dalke, A. & Schulten, K. VMD: Visual molecular dynamics. Journal of Molecular Graphics & Modelling 14, 33–38 (1996). 10.1016/0263-7855(96)00018-5

67 De Lano, W. The PyMOL molecular graphics system. San Carlos CA, USA: De Lano Scientific (2002).

