## Supporting information for "Modulation of *Ideonella sakaiensis* PETase active site flexibility and activity on morphologically distinct substrates by surface charge engineering"

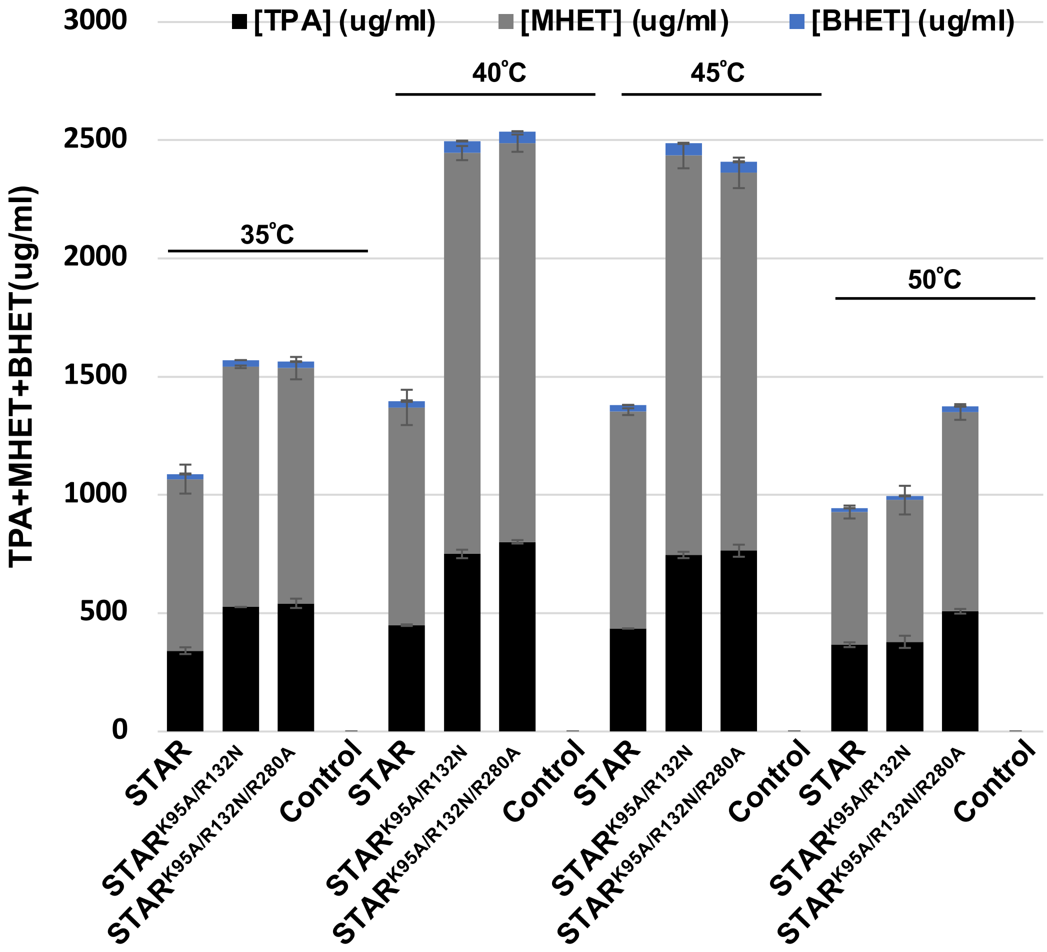


**Supporting Figure 1** PET hydrolysis products generated by indicated enzymes after incubation with lcPET powder substrate for 24 hours. n=3 ± SD.

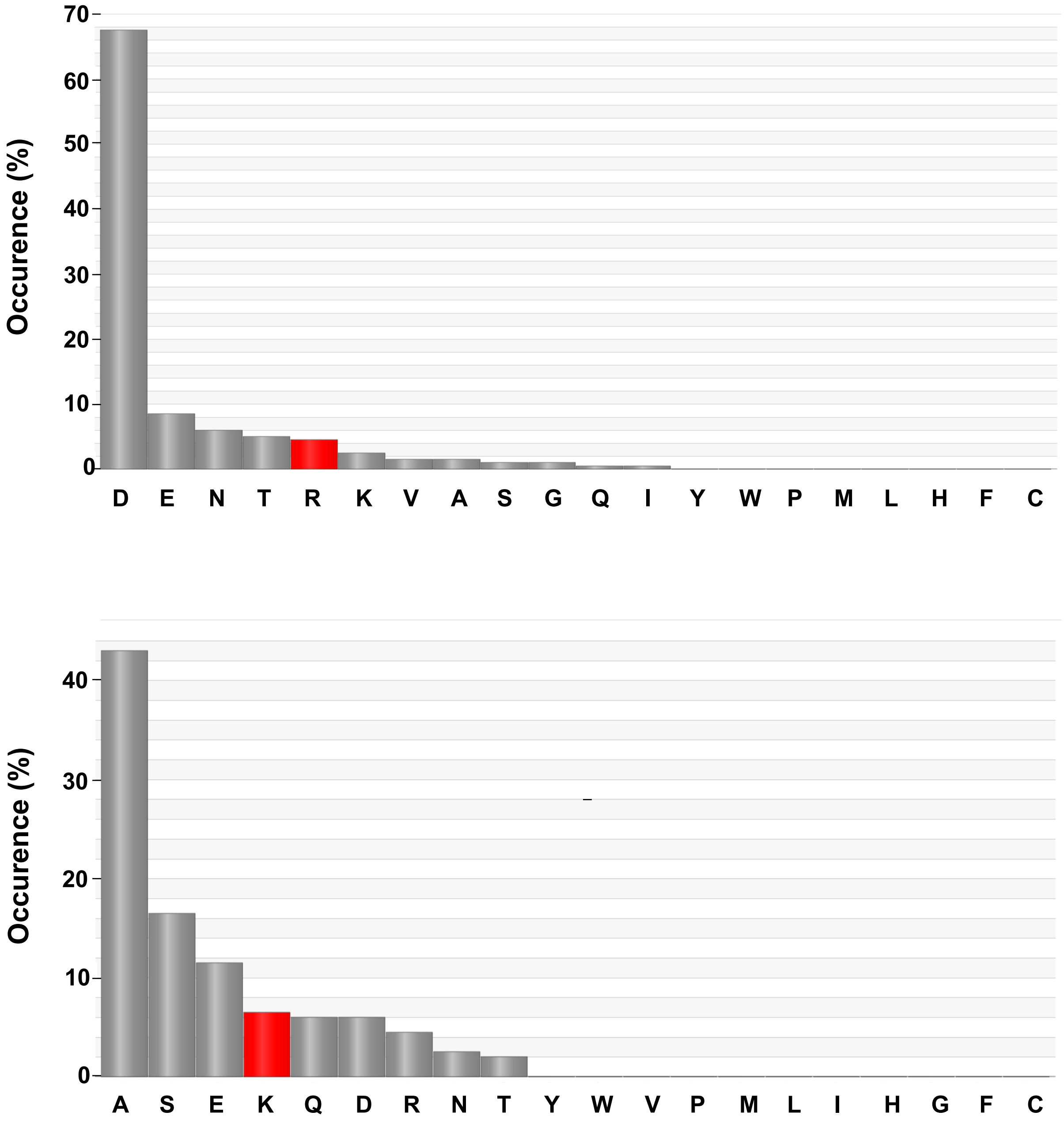


**Supporting Figure 2** Frequency of amino acids present at R132 position (top) and K95 position (bottom) of PETase in 200 evolutionary related proteins. Data generated using HotSpot Wizard.[1]

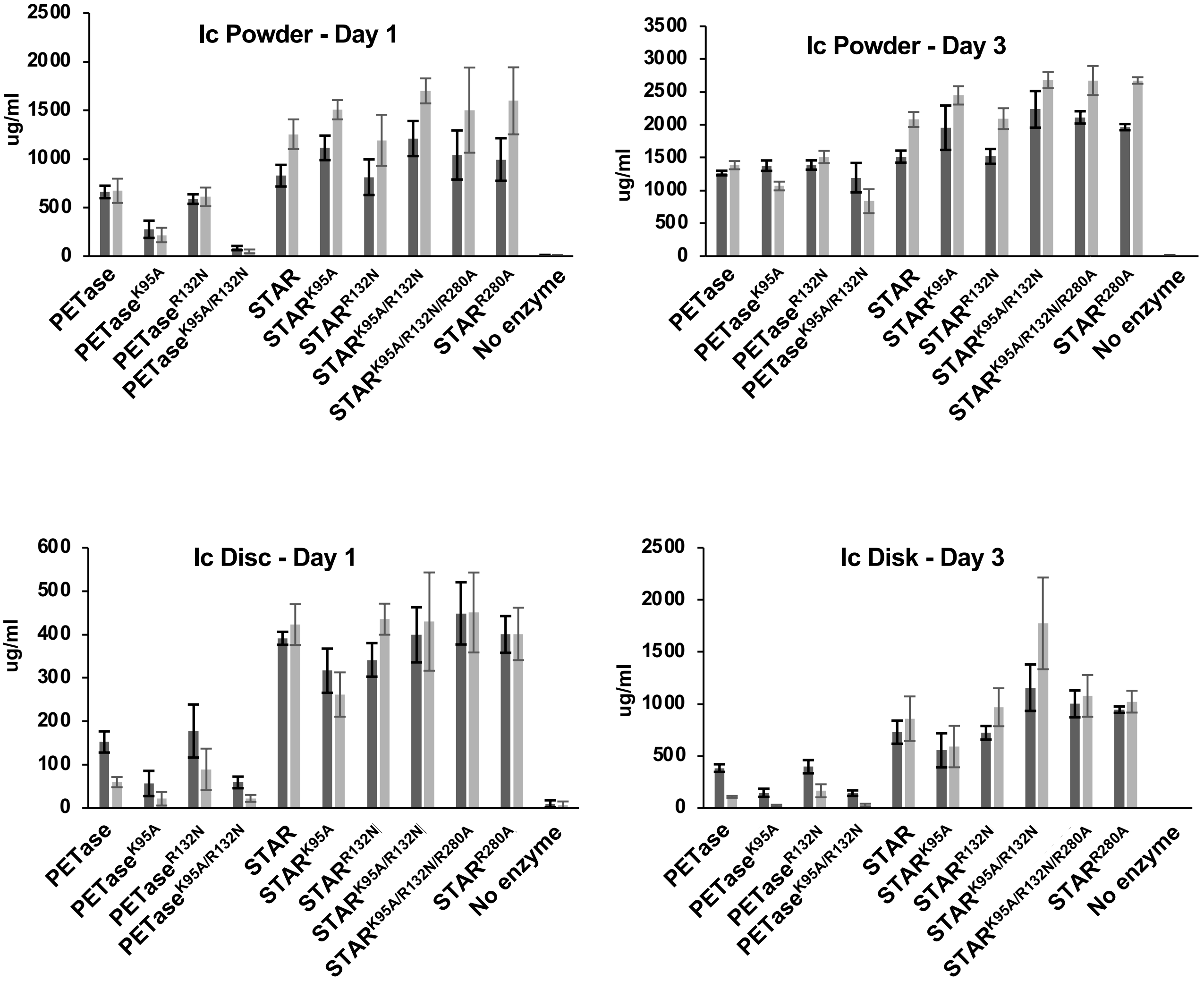


**Supporting Fig 3** TPA (dark grey) and MHET (light grey) hydrolysis products generated by indicated enzymes on Ic powder and film substrates. n=3-5 ± SD.

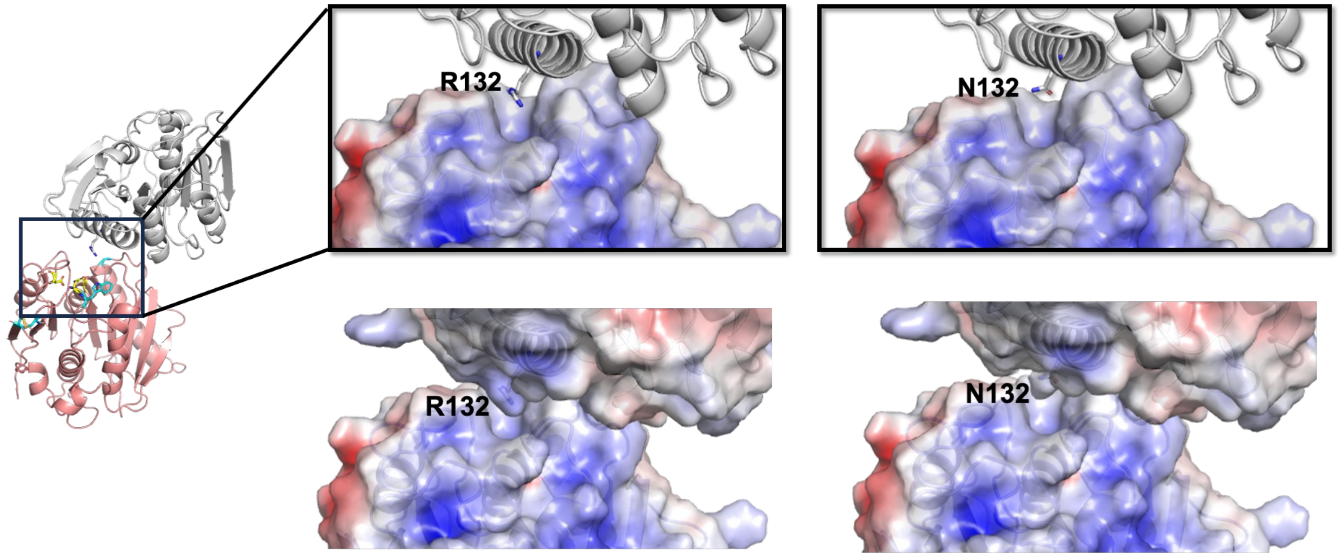


**Supporting Fig 4** Side chain of R132 from PETase monomer projects into substrate binding groove in the vicinity of PETase catalytic triad of another monomer in crystal structure. PETase structures shown in salmon and grey. Active site residues (S160, D206, H237) shown in yellow, STAR-specific residues are shown in cyan. (right) PETase surface electrostatic potential with negative and positive regions depicted red and blue respectively. Adapted from 5XGO.[2]

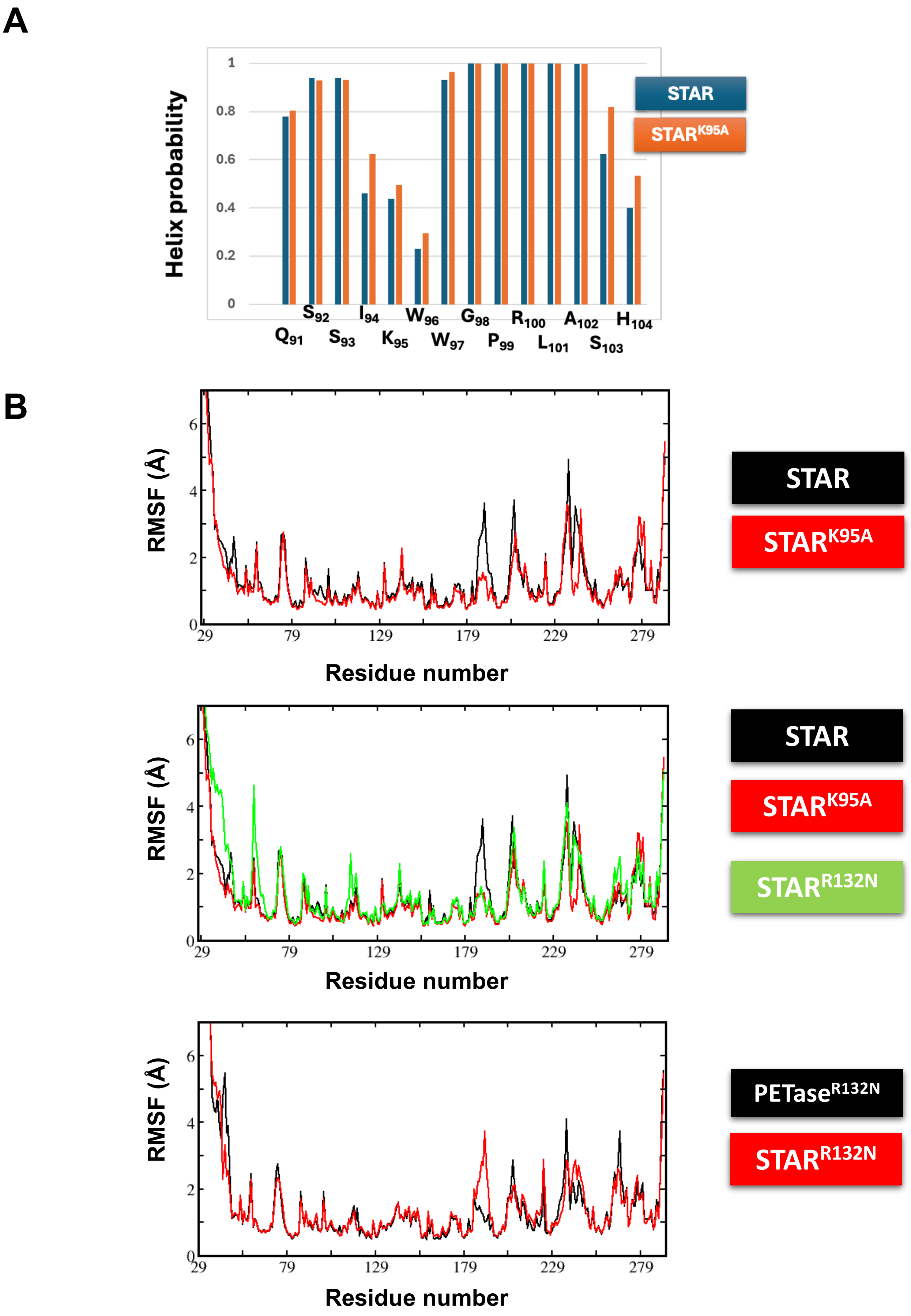


**Supporting Fig 5** K95A and R132N affect enzyme stability and flexibility. A. Helical propensity of indicated residues in STAR and STAR$^{K95A}$ backgrounds calculated by MD simulations**.** B. Root mean square fluctuations (RMSF) of indicated residues in STAR, STAR$^{K95A}$, STAR$^{R132N}$ and PETase$^{R132N}$ enzymes.

### Supporting Figures references

1) Sumbalova, L., Stourac, J., Martinek, T., Bednar, D., Damborsky, J. 2018: HotSpot Wizard 3.0: Web Server for Automated Design of Mutations and Smart Libraries based on Sequence Input Information. *Nucleic Acids Research* 46 (W1): W356-W362 (2018).

2) Han, X., Liu, W., Huang, J.W., Ma, J., Zheng, Y., Ko, T.P., Xu, L., Cheng, Y.S., Chen, C.C., Guo, R.T. Structural insight into catalytic mechanism of PET hydrolase. *Nat Commun* 8: 2106-2106 (2017).
